# Development of a high-throughput protein array to discover selective dsRNA binding domain interactions

**DOI:** 10.64898/2026.11.18.767677

**Authors:** Nicholas Zehrbach, Leilei Shi, Biji Chatterjee, Junwoo Lee, Isaiah Mixon, Mark T Bedford, Charles A. Ishak

## Abstract

Endogenous double stranded RNAs (dsRNAs) function as signal transduction molecules through interactions with host proteins. These interactions can induce signaling cascades implicated in normal homeostasis and diseases such as cancer. Therefore, the discovery of interactions between dsRNAs and dsRNA binding proteins (dsRBPs) is critical for comprehensive elucidation of signal transduction cascades. Discovery of dsRNA-dsRBP interactions remains hindered due to technical limitations that can include a lack of reliable reagents to enrich a given dsRBP or enrichment techniques that cannot identify direct dsRNA-dsRBP binding. As an alternative, we report the development of protein domain arrays to enable high-throughput discovery of direct interactions between dsRNAs and dsRBP ‘reader’ domains. We establish a modular antibody-independent approach coined ‘dsRBP-seq’ to identify putative dsRNAs. Validated dsRNAs are subsequently used to probe dsRBP domain arrays to discover multiple dsRNA binding domains that bind the same dsRNA. The establishment and application of this new technical approach reveals a domain-specific interaction of the dsRNA transcript encoding 26S proteasome complex subunit, Sem1, that binds with double-stranded RNA binding motif 3 of Staufen Double-Stranded RNA Binding Protein 2 (STAU2). Altogether, our workflow establishes a high-throughput approach to discover direct dsRNA-dsRBP interactions and the protein domains required for dsRNA binding.

## INTRODUCTION

Endogenously produced dsRNAs function as signal transduction molecules through interactions with host proteins. In these signal transduction cascades, dsRNA binding domains within dsRNA binding proteins (dsRBPs) function as ‘readers’ to detect and bind dsRNAs (1). Molecular processes that leverage such interactions include mechanisms of RNA decay or innate immune responses (2). Despite their central roles in signal transduction, mechanisms and contexts underlying the employment of specific dsRNA-dsRBP interactions in somatic mammalian cells remain poorly understood. A major contributing factor to this knowledge gap is the current technical limitations that hinder the discovery and characterization of new dsRNA-dsRBP interactions.

Current approaches employed to identify dsRNA-dsRBP interactions have yielded important insights into dsRNA biology. Such widely applied techniques include cross-link immunoprecipitation sequencing (CLIP-seq) and subsequent derivatives of this approach (3). CLIP has been applied to discover putative dsRNA repertoires that bind a given dsRBP. CLIP accuracy depends upon the availability of specific antibodies for a putative dsRBP (3). Therefore, lack of specific antibodies for an endogenous dsRBP limits the utility of CLIP to discover the dsRNA ligands of that protein, and requires tagging the protein of interest to circumvent this limitation. Antibody dependence also hinders the use of CLIP to map minimal interaction domains using dsRBP fragments that lack antibody epitopes. More importantly, CLIP enriches for ribonucleoprotein complexes and therefore does not provide evidence of a direct RNA-protein interaction. Alternatively, antibody-independent approaches have been applied to assess direct dsRNA-dsRBP interactions *in vitro*. For example, RNaseA protection assays have been used to identify dsRNAs that persist after binding to a dsRBP in the presence of RNaseA digestion (4). This approach can reveal domain-level interactions via the use of purified dsRBP fragments. Both CLIP and RNaseA protection assays result in lists of putative dsRNA binders for a given dsRBP. However, acquiring the inverse information regarding dsRBPs that commonly bind a given dsRNA requires the use of alternative approaches.

Current approaches to identify repertoires of dsRBPs that commonly bind a given dsRNA of interest include mass spectrometry following the enrichment of a tagged dsRNA via immunoprecipitation or pulldown (5). However, this approach has largely been applied to discover binders of synthetic dsRNA (6) sequences. As per CLIP, this approach is limited by an inability to identify the precise dsRNA binding ‘reader’ domains across all protein ‘hits’ for a given dsRNA. Altogether, there is an unmet technical need for new approaches that identify new dsRNA-dsRBP interactions with domain-level detail that circumvent the technical limitations of current approaches described above.

In this study, we report the development and application of protein domain arrays to enable high-throughput discovery of direct and specific interactions between dsRNAs and dsRBP reader domains. In doing so, we establish a modular antibody-independent approach coined ‘dsRBP-seq’ to capture putative dsRNAs for subsequent biochemical validation. Validated dsRNAs are then used to probe the dsRBP domain array to discover multiple dsRNA binding domains that bind the same dsRNA target. We apply this technical workflow to discover previously unreported dsRNAs that selectively bind dsRNA binding domain 3 of Staufen Double-Stranded RNA Binding Protein 2 (STAU2). Altogether, our combined workflow of dsRBP-seq with the dsRBP domain array enables high-throughput discovery of new dsRNA-dsRBP interactions across any biological system.

## MATERIALS AND METHODS

### Plasmids

The GST-RNase III WT plasmid was created by inserting the RNase III sequence from the plasmid pRnc-gGFP129, kindly deposited by Marc Ostermeier (7) (189536; Addgene), into the pGEX-4T1 plasmid (s) utilizing cold fusion cloning (MC010B-1; System Biosciences). For the mutant RNase III mutants D45A, E117K, E117A and E117Q, the vector was digested using BamHI-HF (R3136S; New England Biolabs) and SalI-HF (R3138; New England Biolabs), and then point mutations were generated using site directed mutagenesis. All GST fusion plasmids used to express and purify recombinant proteins used in the RBD array were generated as previously described (2).

### Cell culture

Murine ID8 ovarian surface epithelial cells were a generous gift from Dr. Iian McNeish (University of Glasgow, (8)). Adherent cell cultures were grown in Dulbecco’s Modified Eagle Medium (10-013-CV; Sigma), supplemented with 5% (vol/vol) heat-inactivated fetal bovine serum (FBS; 26140-79; Gibco), 1% (vol/vol) of 1.0 g/L Insulin, 0.55 g/L Transferrin, and 0.00067 g/L Selenium (41400045; Gibco), and 10,000 U/mL penicillin with 10,000 µg/mL streptomycin (5% vol/vol) Penicillin-Streptomycin (15140163; Fisher). Cells were maintained at 37°C in 5% CO₂ and routinely screened for mycoplasma using a Mycoplasma PCR Detection Kit (G238; Applied Biological Materials Inc).

### Total RNA extraction for pulldown

For all RNA experiments, a sterile environment was maintained with RNAZap (AM9780; Invitrogen) to destroy RNase A. RNA was isolated using TRIzol according to the manufacturer’s protocol (15596018; Thermo Fisher). To briefly describe, cells were grown on a 15 cm^2^ tissue culture-treated plate (430599; Corning) to approximately 80% confluency. Media was aspirated and 1 mL of TRIzol was added to the plate. Cells were gently scraped and transferred to a 1.7 mL microcentrifuge tube. 0.2 mL of chloroform was added, and the tube was vortexed for 5 seconds. The cells were incubated at room temperature for 5 minutes, and then clarified by centrifugation at 12,000 x g at 4°C for 15 minutes. The aqueous phase was then transferred to a separate 1.7 mL microcentrifuge tube and then incubated with 1 mL 100% isopropanol at 4°C for 10 minutes. The RNA was pelleted by centrifugation at 12,000 x g at 4°C for 15 minutes and then washed 1 mL of 75% ethanol. The tube was then centrifuged at 12,000 x g at 4°C for 5 minutes, and the ethanol was removed. The RNA pellet is air-dried for 10 minutes and then resuspended in 40 µL in RNase-free water. The RNA quantity was evaluated via Nanodrop.

### Recombinant protein expression and purification

GST fusion proteins were generated by transforming plasmids encoding GST-tagged proteins into BL21 (DE3) competent cells according to the manufacturer’s instructions (Thermo Scientific; EC0114). Colonies grown on LB-agar plates in the presence of antibiotic selection using 100 µg/mL ampicillin (A9518; Thermo Fisher) were then selected for inoculation into 20 mL of LB with antibiotic selection 100 µg/mL ampicillin. Cultures were subsequently grown in an incubator shaker (I26; Eppendorf) for 6-8 hours at 37°C shaking at 200 RPM, then spiked with 0.1 mM Isopropyl β-D-1-Thiogalactopyranoside (IPTG, 97063-282; VWR) to induce protein expression for 16 hours at 18°C shaking at 200 RPM. Cells were collected and centrifuged for 10 minutes at 4°C and 5,000 x g. The media was decanted and the cells resuspended in 2 mL of PBS and transferred to 2 mL microcentrifuge tubes. The cells were subjected to sonication using two different pulse codes performed once with a Sonicator (100-132-1639; Fisher Scientific) and converter (102C; Fisher Scientific). Pulse code 1 was set to 16s total, 0.5s interval, and 30% amplitude; Pulse code 2 was set to 10s total, 0.5s interval, and 30% amplitude. Cell contents were clarified by centrifuging the cells at 10,000x g. Protein was then purified by incubating the remaining ∼2mL of supernatant with 50 µL of prewashed 50% glutathione-conjugated agarose bead slurry (16100; Pierce) rotating at 4°C overnight. The beads bound to GST-tagged protein(s) were then spun and washed with 1 mL of 1X PBS at 4°C for 5 min. at 5000 x g. The beads were then transferred to a clean 1.7 mL microcentrifuge tube and washed and spun once more with 1 mL of PBS. The beads were washed and spun a third time with 1mL of wash buffer (100 mM Tris–HCl, pH 8.0, and 120 mM NaCl). Afterwards, the beads were resuspended with 250 µL of elution buffer (100 mM Tris–HCl, pH 8.0, 120 mM NaCl, and 30 mM reduced glutathione) and incubated at 4°C rotating end-to-end for 4 hours. The eluted protein was separated from the beads by centrifugation at 4°C for 15 min and 10,000g. Protein was further purified by dialysis to remove residual bacterial nucleic acids and glutathione: entire protein sample was transferred into SnakeSkin dialysis tubing with 3.5 kDa molecular weight cut-off (Thermo Scientific; PI88242) and subsequently incubated rotating in ≥1 L of 1X PBS for 16 hours at 4°C. After one 1X PBS buffer exchange, protein was further incubated rotating in 1X PBS for 1 hour at 4°C. Protein sample was then transferred from dialysis tubing to a 1.7 mL microcentrifuge tube for protein quantification. Protein quantitation was evaluated using BCA protein assay kit (Thermo Scientific; 23225) as per manufacturer’s protocol, and expected protein size was evaluated utilizing gel electrophoresis (SDS-PAGE).

### Gel electrophoresis and staining

The purified GST protein was normalized and loaded at 0.5 µg per well of a polyacrylamide gel. To prepare for loading, protein samples were diluted to normalize volume, then heated in 10 µL 4X Laemmli Sample Buffer (1610747; Bio-Rad) with 10% 2-Mercaptoethanol (M3148; Sigma Aldrich) at 95°C for 5 minutes. Samples were loaded into TGX Stain-Free Protein Gels (4568084; Bio-Rad) and ran at 80V in a Mini-PROTEAN® Tetra Vertical Electrophoresis Cell (1658004; Bio-Rad) until the loading dye front left the gel. To visualize total protein content, the gel was washed 3X with distilled water and then stained with GelCode Blue (PI24594; Thermo Scientific) for one hour. The gel was then washed with distilled water at room temperature overnight before imaging on a ChemiDoc MP Imaging System (12003154; Bio-Rad).

### GST Pulldown and RNA extraction

GST pulldown was conducted using the general procedure previously described (Ishak *et.* al., 2015) with specific modifications described here. 10 µg of GST protein was bound to 50 µL of prewashed 50% glutathione-conjugated agarose beads by incubating overnight end-to-end at 4°C, in 300 µL of binding buffer (50 mM Tris pH 7.5, 150 mM NaCl, 0.05% NP-40). The bead-protein complex was washed 3 times in 1 mL of binding buffer by spinning for 5000 x g for 5 minutes at 4°C and then removing the supernatant. The washed bead protein complex was resuspended in binding buffer (500 mM NaCl, 200 mM HEPES, 20 mM MgCl_2_, 2 mM dithiothreitol) (4) and incubated with 30 ug of either total RNA or dsRNA generated by *in vitro* transcription (IVT), to a final volume of 250 µL. The reaction was incubated for 1 hour end-to– end at 4°C to promote protein-RNA binding. After the pulldown, 1 mL of TRIzol was added to each reaction and the RNA was isolated, as previously described above, with care to avoid drawing the beads. For the pulldowns, 1 µL of 15 mg/mL GlycoBlue (AM9515; Life Technologies) was added during the ethanol precipitation step to improve visibility of the RNA pellet.

### RNase Digestion assays and RNA extraction

All enzymatic digestions were performed according to the manufacturer’s instructions and suggested conditions. 10 µg of RNA from either isolated RNA or IVT-generated RNA were incubated with RNase If to digest ssRNA (M0243; New England Biolabs), RNase III (M0245; New England Biolabs) to digest dsRNA, or RNase H (M0297; New England Biolabs) to digest RNA:DNA hybrids. All reactions were scaled to a final volume of 20 µL. Afterwards, the reaction was halted: the RNase If reaction through heating at 70°C for 20 minutes, the RNase III reaction through adding 2 µL of 10X EDTA for a final concentration of 1X, and RNase H through adding 1 µL of 0.5M EDTA. Remaining RNA was then isolated from the reaction components through phenol chloroform extraction. The volumes of each reaction were increased to a final volume of 100 µL using nuclease-free distilled water, and then 100 µL of chloroform: phenol isoamyl alcohol (25:24.1; C0549; Sigma Aldrich) was added. The mixture vortexed for 5 seconds and then spun at 12,000 x g for 5 minutes at room temperature. The aqueous phase was transferred to a 1.7 mL microcentrifuge tube, and then incubated with 10 µL of 5 M NaCl, 1 µL of GlycoBlue, and 300 µL of ice-cold 100% ethanol. The tube was mixed by inverting 5 times, then incubated at –20°C for one hour. The RNA was precipitated by centrifugation at 12,000 x g for 5 minutes at room temperature. The pellet was washed once with 70% ethanol, and then air dried for 10 minutes. The RNA was then resuspended in 10 µL of RNase-free water and used immediately for analysis.

### RNA Immunoprecipitation

RNA was isolated through immunoprecipitation using the MAGNA RIP kit (17-700; Sigma Aldrich), according to the manufacturer’s instructions. For each immunoprecipitation, two 15 cm^2^ plates of ID8 cells were grown to 70% confluency. Cells isolated from each plate were lysed in 115 µL of cold RIP Lysis buffer (CS203176, supplemented with the supplied protease inhibitor cocktail (535140, Sigma Aldrich) and RNase inhibitor (CS216144, Sigma Aldrich). Along with 9D5, a supplied IgG control pulldown was used to confirm antibody specificity, using 10 µg of each antibody. RNA quantity was assessed using a Nanodrop, and 5 µL of each condition subjected to RNA dot blot. Isolated RNA was submitted for library preparation and sequencing using the same conditions as per the GST pulldown.

### *In vitro* transcription

Sem1 and Pfn1 were selected for *in vitro* validation from the genes enriched in the GST–RNase III E117A pulldown relative to the matched input, as defined in the Sequencing analysis section. Reference mRNA sequences were retrieved from the NCBI Nucleotide database (https://www.ncbi.nlm.nih.gov/nuccore) using the RefSeq transcript for each gene (Sem1, NM_009169.2, 491 bp; Pfn1, NM_011072.4, 883 bp). Gene-specific primers were designed against these sequences with Primer-BLAST (https://www.ncbi.nlm.nih.gov/tools/primer-blast/). Primer melting temperature (Tm) was constrained to a minimum of 55 °C, an optimum of 59 °C and a maximum of 63 °C, with a maximum Tm difference of 3 °C within a pair. The maximum PCR product size was set to the full length of the corresponding transcript (491 bp for Sem1, 883 bp for Pfn1) so that candidate amplicons could span as much of the mature mRNA as possible. Primer-pair specificity was checked against the RefSeq mRNA database for genic targets and the RefSeq representative genome database for intergenic regions, with the search organism restricted to Mus musculus (taxid: 10090); all other parameters were left at their default values.

Three forward–reverse primer sets were designed per gene, and pairs predicted to amplify off-target transcripts or genomic loci were excluded. For each gene, the highest-ranked pair returned by Primer-BLAST was selected among those with a predicted amplicon longer than 300 bp **(Supplemental Table S2)**. This length criterion was applied because transcripts shorter than 0.3 kb require a modified in vitro transcription set-up with increased template and extended incubation to reach maximum yield (NEB #E2040 protocol); restricting candidates to longer amplicons allowed all templates to be transcribed under identical standard conditions.

For each selected pair, the T7 promoter sequence (5′-TAATACGACTCACTATAGGG-3′) was appended to the 5′ end of the forward primer only, so that PCR amplification yields a linear double-stranded template directing synthesis of the sense-strand RNA alone. Antisense transcription was not performed: inspection of the dsRBP-seq coverage in IGV showed no evidence of divergent or antisense transcription at either locus, indicating that the double-stranded character recognized by RNase III E117A is more likely to arise from intramolecular folding of the single-stranded mRNA than from sense–antisense duplex formation.

Total RNA isolated from ID8 cells was used to prepare IVT products. 1 µg of RNA was converted into cDNA by using the SuperScript IV VILO Master Mix with ezDNase enzyme (11766050, ThermoFisher), according to the manufacturer’s instructions. The cDNA was quantitated via Nanodrop, and then 10 ng of cDNA was amplified using the Q5 High-Fidelity Master Mix (M0492, NEB) according to the manufacturer’s instructions. 10 µM of forward primer and 10 µM of reverse primer were used in each experiment. The PCR reaction was performed with the following conditions in a thermocycler (1861096, Bio-Rad): initial denaturing step at 98°C, for 30s. Amplification over 27 cycles at 98°C for 10s, then at 60°C for 20s, and 72°C for 20s. The final extension is set to 72°C for 2 min. After PCR, 0.5 µg DNA is loaded in a 0.5% w/vol Tris-Boric Acid-Ethylenediaminetetraacetic acid (TBE; 18-136, Apex Bioresearch) gel and ran to separate fragments. DNA Bands are stained using SYBR Safe DNA gel stain (S33102; Invitrogen) according to the manufacturer’s instructions, and the bands for the transcripts of interest are cut under UV light. The gel bands are processed using the NucleoSpin Gel and PCR Clean-up Kit (740609.250, Macherey-Nagel), and multiple wells are pooled together. 1 µG of template DNA is converted to RNA using the (P1320, Promega), according to the manufacturer’s instructions.

### 3’ End Biotin Labeling

Biotin was conjugated to the dsRNA probes using a modified protocol(9). 50 pmol of IVT product was biotinylated using a 3’ End Biotinylation Kit (Pierce, 20160). RNA was heated at 85°C for 5 minutes before proceeding with the reaction. The biotinylation reaction was allowed to proceed for 16 hours at 16°C. After biotinylation, the RNA was purified from the components by using TRIzol as previously described. After purification, the RNA was validated by running 1 µg on a denaturing TBE-Urea gel (4566033, Bio-Rad) to confirm the appropriate size, and by performing a dot blot with α-dsRNA antibody to confirm secondary structure.

### Dot blot and antibodies

RNA dot blots were conducted as previously described (Ishak et. al., 2025) to evaluate the presence of dsRNA. Briefly, samples were pipetted onto a positively charged ‘Hybond-N+’ nylon membrane (RPN303B; Amersham Pharmacia Biotech) at 3 µL per sample. The membrane was left to dry for approximately 15 minutes at room temperature. The membrane was then UV-crosslinked with a UV Crosslinker (FB-UVXl-1000; Fisher Scientific). The membrane was then blocked for 1 hour at room temperature in membrane blocking buffer, 5% milk vol/vol (B08H9ZTRGL; Swiss Miss) diluted in 0.01% PBS/T (PBS/Tween-20, P9416; Sigma Aldrich) vol/vol. The membrane was then incubated overnight at 4°C using either α-dsRNA clone J2 at 1:500 (76651L; Cell Signaling Technologies) or 9D5 (Ab00458-23.0; Absolute Antibody) at 1:1000 in blocking buffer. Afterwards, the membrane was washed 6 times in PBS-T every 5 minutes. The membrane was then incubated for 1 hour at room temperature with the appropriate horseradish peroxidase–conjugated rabbit secondary antibody (7074S; Cell Signaling Technologies) or mouse secondary antibody (7076S; Cell Signaling Technologies), diluted in membrane blocking buffer. The membrane was washed 6 times in PBS-T every 5 minutes before development. The membrane was developed with Femto substrate as per manufacture’s protocol (34096; Life Technologies) and imaged on an Amersham Image Quant 800 (29399481; Cytiva). To evaluate total nucleic acid loading, the membrane was first washed 3 times in PBS-T every 5 minutes to remove residual chemiluminescence reagent. The membrane was incubated for 2 hours at room temperature with 0.5% methylene blue (M9140-25G; Sigma) vol/vol in 30% ethanol (vol/vol). Finally, the membrane was washed 2 times in distilled water and imaged on a ChemiDoc MP Imaging System (12003154; Bio-Rad).

### Library preparation and sequencing

Samples were submitted to Admera Health for library preparation and sequencing. Libraries were prepared using the SMARTer Stranded Total RNA-Seq Kit v2 Pico Input (634411; Takara), according to the manufacturer’s instructions. Quality was assessed using the high-sensitivity (hs)RNA ScreenTape (5067-5579; Agilent technologies) and an Agilent Tapestation. Libraries were sequenced to obtain ≥60M 150 bp paired-end reads per sample using an Illumina NovaSeq X (20084803; Illumina).

### Sequencing analysis

Raw read quality was assessed with FastQC v0.12.1(10) and summarized across samples with MultiQC v1.25.2(11). Adapter sequences and low-quality bases were removed with Trim Galore v0.6.10(12) in paired-end mode with default parameters, and trimmed reads were re-assessed with FastQC and MultiQC.

Trimmed reads were aligned with STAR v2.7.11b(13) to the mouse genome (GRCm38.p6 primary assembly, GENCODE vM25 annotation; –-sjdbOverhang 149). Both datasets were aligned with –-outFilterMultimapNmax 100, –-winAnchorMultimapNmax 100, –-outFilterMismatchNoverReadLmax 0.04, –-alignIntronMin 20, –-alignIntronMax 1000000, –– alignMatesGapMax 1000000, –-alignSJoverhangMin 8 and –-alignSJDBoverhangMin 1, with all other parameters at default unless stated below.

The 9D5 RIP-seq libraries were aligned to the mouse genome alone, with –– seedSearchStartLmax 30 and –-outSAMmultNmax –1 (all alignments reported). The dsRBP-seq libraries were aligned to a combined reference of the mouse genome and the Escherichia coli K-12 MG1655 (NC_000913.3) and BL21(DE3) (NC_012971.2) genomes as decoy sequences, with a mouse-only gene annotation. Alignment settings for dsRBP-seq were –-seedSearchStartLmax 20, –-outFilterScoreMinOverLread 0, –-outFilterMatchNminOverLread 0, – –outFilterMatchNmin 30, –-peOverlapNbasesMin 10, –-peOverlapMMp 0.01 and –– outSAMmultNmax 1.

Alignments to non-canonical scaffolds (and, for dsRBP-seq, to the E. coli decoy contigs) were removed with samtools v1.21(14), retaining chr1–chr19, chrX, chrY and chrM. Read groups were assigned with Picard v3.4.0 AddOrReplaceReadGroups, and PCR duplicates were marked, but not removed, with Picard MarkDuplicates. Secondary and supplementary alignments were then removed (samtools view –F 0×900), so only primary alignments were used downstream. No mapping-quality filter was applied.

Strand-specific coverage tracks were generated with deepTools v3.5.6 bamCoverage(15) using –-extendReads, –-binSize 10 and –-filterRNAstrand forward or reverse. Each track was normalized to counts per million (CPM), computed separately for each strand-filtered track. Tracks were visualized in the Integrative Genomics Viewer(16).

Gene-level enrichment analysis was performed for the dsRBP-seq dataset only. Gene-level read counts were generated by STAR (--quantMode GeneCounts) against the GENCODE vM25 annotation; these counts include uniquely mapped reads only. Because the libraries are reverse-stranded, the reverse-strand count column was used. Analysis was restricted to protein-coding genes, and Lars2 was excluded because its locus absorbs rRNA-derived reads.

Enrichment of RNase III E117A pulldown over input was tested in edgeR v4.4.3(17). Lowly expressed genes were removed with filterByExpr, libraries were normalized with the TMMwsp method, and differential enrichment was tested with exactTest. Each contrast had one library per condition, so dispersion could not be estimated from the data. It was therefore fixed at a biological coefficient of variation (BCV) of 0.2 (dispersion = 0.04), following edgeR guidance for genetically homogeneous cell-line material; the resulting p-values are conditional on this assumed dispersion. P-values were adjusted with the Benjamini–Hochberg method. Genes were classified as enriched or depleted at FDR < 0.05 and |log2 fold change| ≥ 1 in the RNase III E117A-versus-input contrast.

### Protein array

The dsRNA-binding domain–focused protein arrays were generated and probed as previously described (2). The version 1 array comprised five blocks (A–E) containing 51 GST-fusion proteins harboring DSRM, DND1, helicase, RIG-I C-terminal, LRR, DSRI, and OB-fold domains. The version 2 array included all proteins from version 1, along with an additional block (F) containing full-length MDA5, RIG-I, LGP2, TLR3, RNase III, and RNase H, as well as mutant forms of MDA5 and RNase III. The biotinylated poly-A was synthesized by Integrated DNA Technologies (IDT), and biotinylated polyI:C was purchased from Sigma (P1530-25MG). For Sem1 and Pfn1, RNA was generated by in vitro transcription and biotinylated as described above. Briefly, biotinylated probes were incubated with streptavidin–Cy3 (Cytiva, catalog no. PA43001) for 30 minutes at 4°C, followed by the incubation with Biotin-Agarose beads (Sigma, B0519-5ML) to remove unbound streptavidin–Cy3. The Cy3 labeled probes were then incubated with the array slides overnight. The following day, the slides were washed three times with RNA washing buffer (20 mM Tris–HCl, pH 7.4, 200 mM KCl, 0.2 mM ethylenediaminetetraacetic acid (EDTA), 0.05% NP-40, and 0.4 U RNase inhibitor) to remove unbound probes. After drying, the slides were scanned using an InnoScan 1100 AL microarray scanner to detect probe-binding signals.

## RESULTS

### dsRBD arrays distinguish dsRNA versus ssRNA substrates

Protein arrays have enabled the discovery of protein ‘reader’ domains that specifically bind modified peptide targets (18). Beyond mapping protein-protein interactions, our recent work demonstrates that protein arrays can also be used to discover protein-RNA interactions (2). Therefore, we sought to determine whether we could extend the utility of domain arrays to specifically capture and identify new dsRNA-protein interactions. To generate dsRNA-domain binding arrays, we selected domains to array from our previously published RNA recognition motif (RRM) array (2). Candidates were selected based upon the presence of putative dsRNA binding domains (2). Following selection, domains were expressed with an N-terminal GST tag in bacteria, and purified as we previously describe (2,18). GST-tagged domains were then arrayed in separately positioned technical duplicate spots onto a nitrocellulose-coated glass slide with ‘GST-only’ negative controls arrayed at the center of every protein block as previously described (2).

To test the specificity of our putative dsRNA-domain binding array, we conducted our previously optimized workflow in which biotinylated RNA probes are bound to separate arrays (2). Biotinylated RNA probes are first labeled with Cy3-conjugated streptavidin, and unbound streptavidin–Cy3 is removed using biotin–agarose beads. The labeled probes are then incubated with the array. Following washing to remove unbound probes, fluorescence is detected as an indicator of RNA binding to the arrayed peptides and proteins **(Figure 1A)**. Fluorescence signal is then mapped back to the array map to identify domains that produce specific signal in technical duplicates for a given RNA probe **(Figure 1B)**. Using this workflow, we tested array specificity by comparing binding profiles of a biotinylated synthetic ssRNA probe (poly(A)) versus a biotinylated synthetic dsRNA mimetic (poly(I:C)) (2). We observe strong fluorescence signal on the array following the use of poly(I:C) but not poly(A) **(Figure 1C)**. Consistent positive signal across technical duplicates of each positive ‘hit’ domain combined with lack of positive signal at the GST-only control spot confirms signal specificity. Therefore, these results validate that our dsRNA-domain binding array can specifically detect protein interactions to dsRNAs.

**Figure 1:**
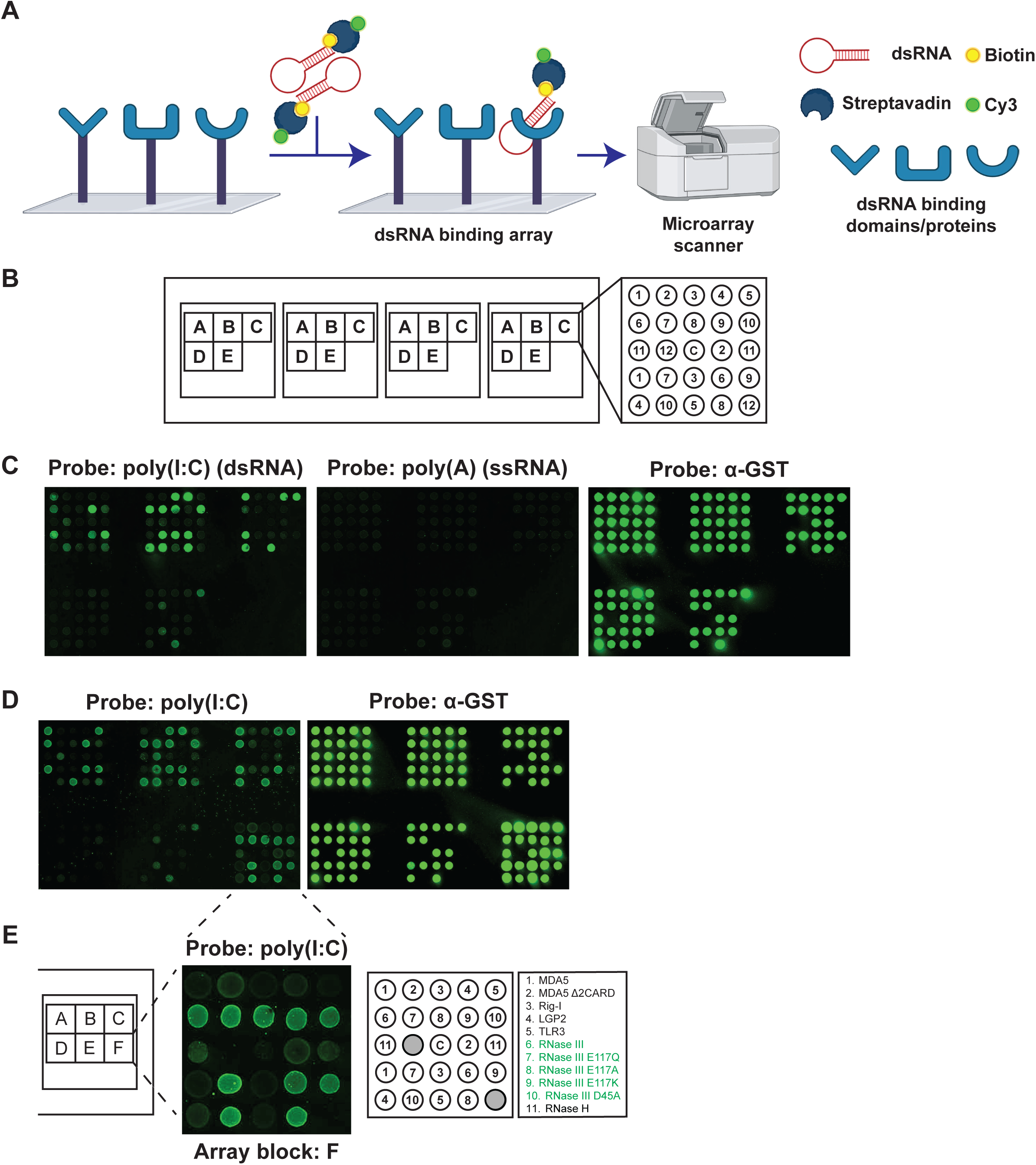
Development and validation of domain arrays to capture dsRBP-dsRNAs interactions. **(A)** Representative schematic of dsRBP array. Domains/proteins are printed onto nitrocellulose glass slides. The different biotinylated RNA probes labeled with Cy3-streptavidin are then incubated with the slides. After washout of unbound probes, slides are then imaged to detect fluorescence signal. The schematic was generated using BioRender. **(B)** Layout of proteins on slides. Proteins were printed on the slide, and each protein has two distinct locations orientated based on block (A-E), with one ‘GST-only’ loading control in the center of each block labelled “C”. **(C)** Representative fluorescence image from array version 1 of either Cy3-detected probe of the dsRNA mimetic poly(I:C), the ssRNA mimetic poly(A) or of fluorescence signal from α-GST antibody for loading. **(D)** Representative fluorescence image from array version 2 of either Cy3-detected probe of the dsRNA mimetic poly(I:C), or of fluorescence signal from α-GST antibody for loading. **(E)** Representative layout of block F from array version 2. **(F)** Representative inset of block F, highlighting the signal from poly(I:C). Full-length proteins incorporated on the block are listed with positive ‘hits’ highlighted based on signal from both technical replicates.

Finally, we sought to expand our dsRNA-domain binding array to include full length proteins not originally included on our previously published RNA recognition motif (RRM) array (2). Additional candidates selected include the dsRNA-specific nuclease RNase III, previously characterized catalytically inactive RNase III mutants (19–23), along with proteins that sense cytosolic dsRNAs such as members of the RIG-I-like family of proteins and TLR3 **(Supplementary Figure S1A)**. Repetition of binding using biotinylated poly(I:C) produced positive signal at the newly added dsRBPs but not at the GST-only control **(Figure 1D-E)**. Altogether, our results reveal that arrayed dsRBP domains retain the ability to bind dsRNAs and distinguish RNA substrates based upon secondary structure.

### GST pulldowns using recombinant dsRBPs specifically enrich dsRNAs *in vitro*

We sought to test whether the dsRNA-domain binding array could be used to discover multiple dsRBPs that bind the same dsRNA substrate. To do so, we strategized to test dsRNAs identified as binders to at least one of the domains on the array. We selected RNase III as a broad dsRNA binding protein to establish proof-of-principle. To identify dsRNA binders to RNase III, we first tested the ability of purified GST-RNase III **(Figure 2A)** to specifically bind dsRNAs following a pulldown with total RNA. In parallel, we included the catalytically inactive RNase III^E117A^ mutant **(Figure 2B)** to enable downstream sequencing while minimizing the risk of digesting captured RNAs. Following protein purification **(Figure 2C)**, we conducted GST-pulldowns using GST-RNase III, GST-RNase III^E117A^, and a GST negative control with total RNA as input **(Figure 2D)**. Input RNA was isolated from murine ID8 cells that were previously shown to express dsRNAs (24). RNA bound to the GST-tagged proteins was subsequently isolated and subjected to dot blots to assess dsRNA enrichment using the J2 antibody that binds dsRNA (25). We observe strong dsRNA enrichment following pulldowns with GST-RNase III and GST-RNase III^E117A^, but not GST alone. These results demonstrate that GST pulldowns using dsRBPs can be used to isolate dsRNA from total RNA input **(Figure 2E)**.

**Figure 2:**
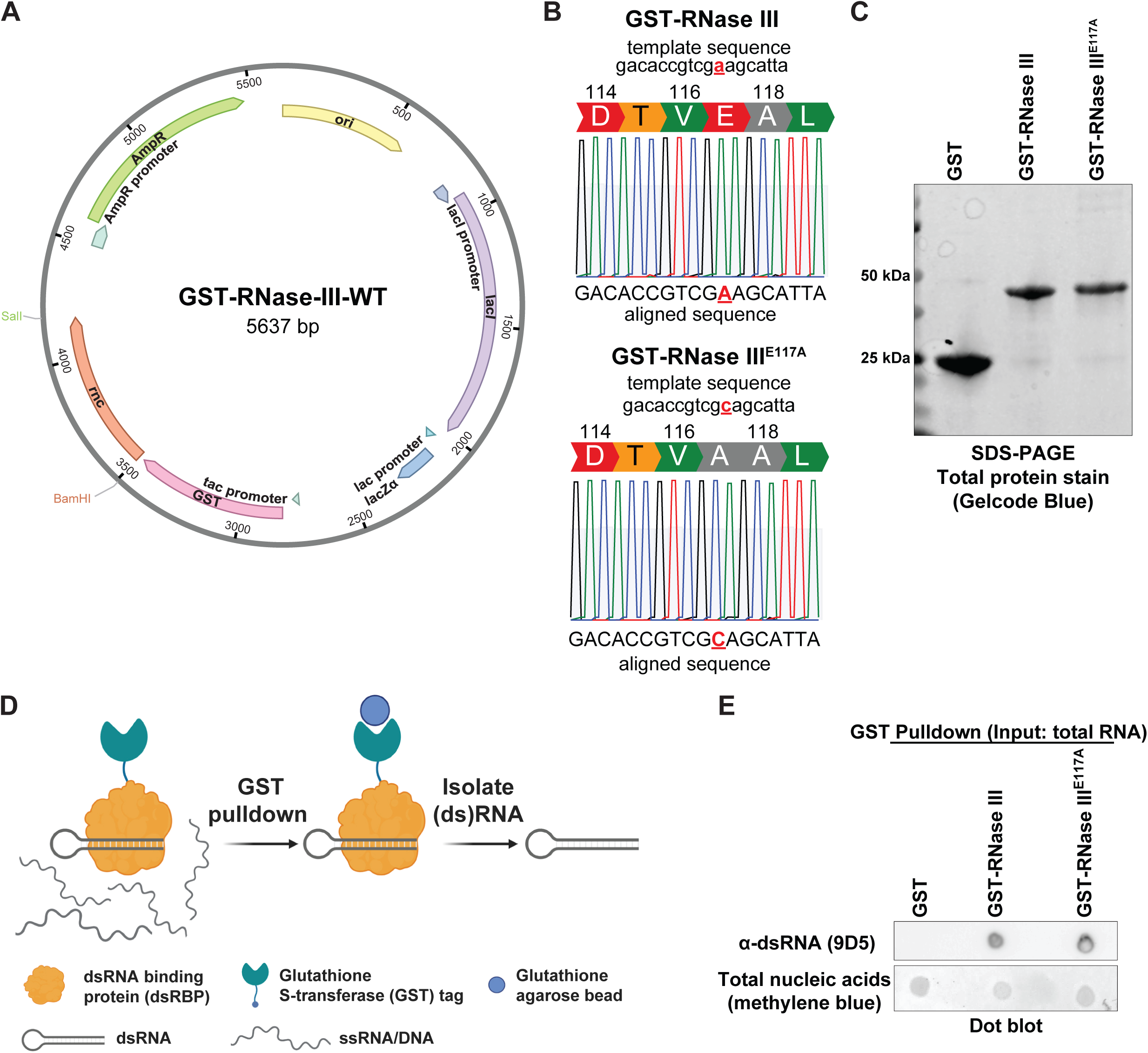
Development of assay to enrich dsRBP-bound dsRNA from total RNA in vitro. **(A)** Plasmid map of full-length RNase III, highlighting plasmid design. **(B)** DNA sequencing chromatograms of plasmids encoding GST-RNase III wild type, and GST RNase III E117A. The mutation is highlighted in red. **(C)** Representative SDS-PAGE gel of purified GST proteins used in pulldown. Proteins were harvested from bacteria, purified, loaded at approximately 0.5 ug and stained for protein using GelCode Blue. **(D)** Schematic of GST-dsRBP pulldown workflow. Purified GST-proteins were incubated with GSH-beads and then incubated with total RNA from ID8 cells. dsRNA-protein complexes were isolated from total RNA via centrifugation and wash steps, then purified via TRIzol isolation. RNA was subject to dot blot. The schematic was generated using BioRender. **(E)** Representative dot blot illustrating protein-specific isolation of double stranded RNA. RNA purified after pulldown was used in the dot blot, probed first for dsRNA (clone 9D5), and then stained methylene blue to visualize total nucleic acids.

### dsRBP-seq identifies bona fide dsRNA binding partners of RNase III^E117A^

To identify dsRNAs that bind RNase III and other dsRBPs on our array, we repeated the GST-pulldown using GST-RNase III^E117A^ with total RNA isolated from ID8 cells as input. Isolated RNA was subsequently subjected to library preparation without rRNA depletion to identify all bound candidates, followed by paired-end stranded sequencing to identify candidate dsRNAs bound to RNase III^E117A^ **(Figure 3A)**. To validate the specificity of our assay to enrich dsRNAs, we identified putative enriched dsRNAs that would be amendable to in vitro transcription (IVT) for subsequent biochemical experiments. We prioritized dsRNAs with a length of <1kb with signal entirely derived from exons to enable IVT from reverse transcribed total RNA, and to adhere to recommended oligo size limits for successful 3’ specific biotinylation. Using these criteria, we identified two putative RNase III^E117A^ binding partners for downstream validation: i) *Sem1* **(Figure 3A-B)** and ii) *Pfn1* **(Supplementary Figure S2A)**.

**Figure 3:**
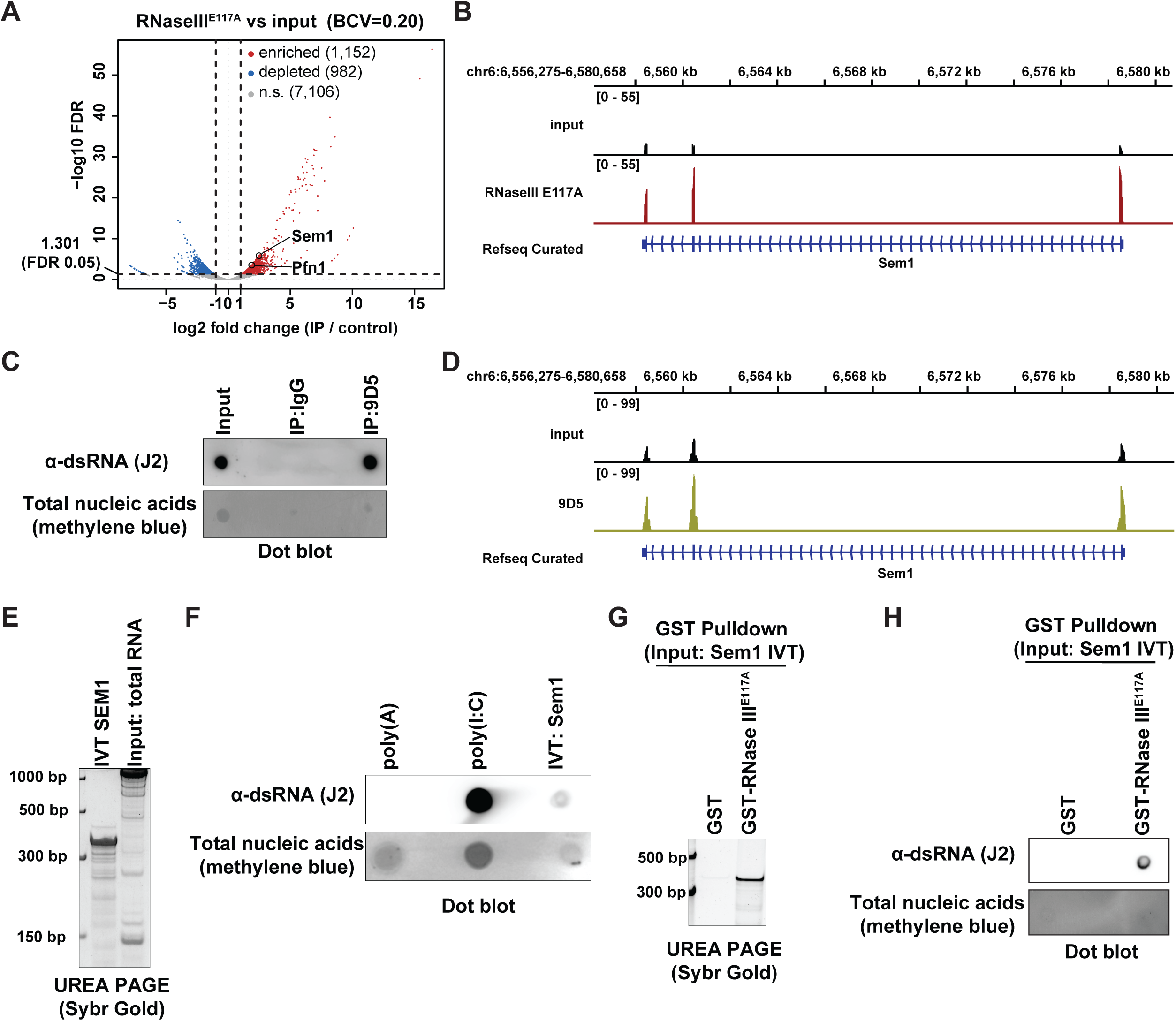
GST-RNase III pulldown sequencing identifies candidate dsRNAs. **(A)** Volcano plot showing enriched and depleted genes from the dsRBP-seq comparing the GST-RNase III E117A pulldown relative to total input RNA. **(B)** GST-RNase III^E117A^ dsRBP-seq signal enrichment at the *Sem1* locus. **(C)** Representative dot blot demonstrating the specificity of the 9D5 antibody to immunoprecipitate dsRNA. RIP was performed using total RNA from ID8 cells immunoprecipitated using either IgG or the dsRNA 9D5 antibody. The presence of dsRNA was assessed using the mouse monoclonal J2 dsRNA antibody. Methylene blue staining shows total nucleic acids. **(D)** 9D5 RIP-seq signal enrichment at the *Sem1* locus. **(E)** Representative denaturing TBE-Urea gel of the in vitro transcribed (IVT) Sem1 candidate sequence, or total RNA. **(F)** Representative dot blot of poly(A), poly(I:C), and IVT Sem1 candidate RNA probed with anti-dsRNA (J2). Methylene blue staining shows total nucleic acids. **(G)** Representative denaturing TBE-Urea gel of the GST-dsRBP pulldown using IVT Sem1. **(H)** Representative dot blot showing dsRNA isolated after the GST-dsRBP pulldown.

We conducted orthogonal assessments to test whether *Sem1* and *Pfn1* identified from dsRBP-seq using GST-RNase III^E117A^ form dsRNAs. First, we conducted RNA immunoprecipitation sequencing (RIP-seq) as previously described (26) to enrich total dsRNA from murine ID8 ovarian surface epithelial cells. Since the commonly used J2 dsRNA antibody is raised in mice (27), we instead used the recombinant 9D5 dsRNA antibody raised in rabbit (28). RNA immunoprecipitation using 9D5 followed by dot blot analysis using J2 confirms that our immunoprecipitation protocol using 9D5 specifically enriches for dsRNA **(Figure 3C)**. Next, we conducted RIP-seq using the same experimental conditions. As observed using dsRBP-seq with GST-RNase III^E117A^, RIP-seq using 9D5 identifies both *Sem1* **(Figure 3C-D)** and *Pfn1* **(Supplementary Figure S2B)** as putative dsRNAs.

With two sequencing-based approaches identifying *Sem1* and *Pfn1* as putative dsRNAs, we proceeded with biochemical validation to test dsRNA formation. We conducted *in vitro* transcription from total RNA isolated from ID8 cells to amplify full length *Sem1* and *Pfn1* RNA.

Resolving the transcription reactions by Urea-PAGE reveals enrichment of prominent bands at the expected sizes of full length 345 bp and 728 bp for *Sem1* **(Figure 3E)** and *Pfn1* **(Supplementary Figure S2C)** respectively. Next, we conducted dot blot analysis using the IVT products to test their ability to form dsRNAs *in vitro*. Dot blots probed with J2 reveal positive signal for *in vitro* transcribed *Sem1* **(Figure 3F)** and *Pfn1* **(Supplementary Figure S2D)**. Collectively, these results establish that *Sem1* and *Pfn1* form dsRNAs *in vitro*.

Next, we sought to validate our dsRBP-seq results by testing the binding of purified GST-RNase III^E117A^ to *in vitro* transcribed *Sem1* and *Pfn1* dsRNA. We conducted GST pulldowns using GST-RNase III^E117A^ and a GST control with *in vitro* transcribed *Sem1* or *Pfn1* dsRNA as the input. RNA purified post-pulldown was resolved by Urea-PAGE, and revealed a prominent band approximately at the expected size of 345 bp for *Sem1* enriched in the GST-RNase III^E117A^ pulldown **(Figure 3G)**. Dot blots of the same RNA probed with 9D5 reveal positive signal specifically in the GST-RNase III^E117A^ pulldown **(Figure 3H)**. Likewise, dot blots reveal that GST-RNase III^E117A^ binds *in vitro* transcribed *Pfn1* **(Supplementary Figure S2E).** Altogether, our biochemical experiments validate our dsRBP-seq results, and demonstrate that *Sem1* and *Pfn1* form dsRNAs that are bound by RNase III^E117A^.

### dsRBP array identifies domain-specific binding of *Sem1* and *Pfn1* to STAU2

Identification and validation of *Sem1* and *Pfn1* as dsRNAs that bind RNase III positioned us to test the ability of our dsRBP array to identify *de novo* binders of *Sem1* and *Pfn1*. To identify candidate domains that bind *Sem1* and *Pfn1* dsRNAs, we 3’ biotin-conjugated *in vitro* transcribed Sem*1* and Pfn1 and subjected each biotinylated dsRNA probe to our dsRBP array workflow **(Figure 1A)**.

The *Sem1* dsRNA probe produced strong fluorescence signal corresponding to specific domains arrayed within blocks B, C, and F **(Figure 4A)**. We confirm that the biotin-conjugated *Sem1* dsRNA probe bound full-length RNase III^E117A^, in addition to wildtype RNase III and the other RNase III mutants within the array **(Figure 4B)**. Surprisingly, we found that certain domains representing distinct fragments of the same protein exhibit different binding affinities for *Sem1* dsRNA. We found that *Sem1* bound strongly with the STAU2 DSRM 3 domain that encompasses Staufen 2 (STAU2) double-stranded RNA binding motif (DSRM) 3. In contrast, *Sem1* did not exhibit the same binding towards STAU2 DSRM 4 or STAU2 DSRM (1–2) domains **(Figure 4C)**.

**Figure 4:**
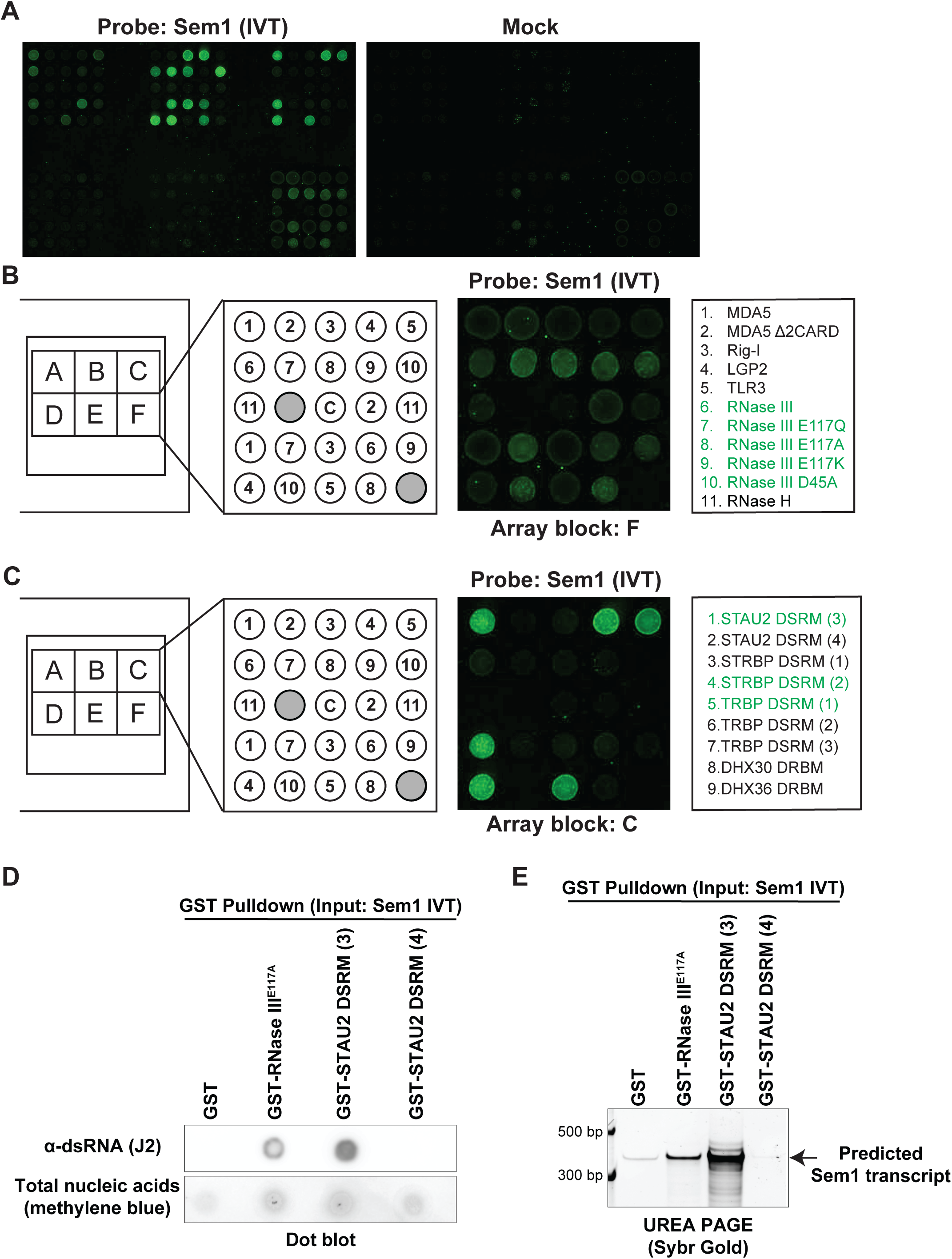
High throughput discovery of dsRNA binding domains that bind Sem1 dsRNA. **(A)** Representative fluorescence image from array version 2 of Cy3-detected probe of Sem 1 and mock treatment. **(B)** Representative inset of block F, highlighting the signal from Sem1. Full-length proteins incorporated on the block are listed with positive ‘hits’ highlighted based on signal from both technical replicates. **(C)** Representative inset of block C, highlighting the signal from Sem1. Domains are listed with positive ‘hits’ highlighted based on signal from both technical replicates. **(D)** Representative blot showing dsRNA isolated after the GST-dsRBP pulldown using (IVT) Sem1. dsRNA signal is displayed, along with methylene blue staining of total nucleic acids. **(E)** Representative denaturing TBE-Urea gel of the GST-dsRBP pulldown using in vitro transcribed (IVT) Sem1.

To validate the results of the array, we performed GST pulldowns using bacterially purified GST-tagged STAU2 DSRM 3 and 4 domains **(Supplementary Figure S3A-B)** with *in vitro* transcribed *Sem1* as the input. After pulldown and RNA purification, dot blots using anti-dsRNA (J2) reveal dsRNA enrichment from pulldowns using GST RNase III^E117A^ and STAU2 DSRM 3 domain, but not the STAU2 DSRM 4 domain **(Figure 4D)**. Resolving the pulldowns via UREA-PAGE confirmed specific binding to *Sem1* dsRNA **(Figure 4E)**. Next, we assessed if discovery of *de novo* protein binders using the dsRBP array could be repeated for *Pfn1* dsRNA. We repeated the array workflow **(Figure 1A)** using in vitro transcribed and biotinylated *Pfn1* dsRNA and observed significant but weaker binding in blocks C and D **(Supplementary Figure S3C-D)**. Similar to *Sem1*, we found preferential differential binding of *Pfn1* dsRNA to STAU2 DSRM 3 versus the STAU2 DSRM 4 domain **(Supplementary Figure S3C-D)**. We then conducted a GST pulldown using GST-tagged RNase III^E117A^, STAU2 DSRM 3, and STAU2 DSRM 4 with *in vitro* transcribed *Pfn1* as the input. Dot blots using anti-dsRNA (J2) of the RNA purified post pulldown reveals that GST-RNase III^E117A^ and STAU2 DSRM 3 bound *Pfn1* dsRNA while STAU2 DSRM 4 does not exhibit binding **(Supplementary Figure S3E)**. Altogether, these data validate the ability of our dsRBP array to uncover *de novo* protein binding partners of a given dsRNA with domain-level specificity.

## DISCUSSION

Interactions between endogenous dsRNAs and host dsRNA-binding proteins contribute towards signal transduction cascades that govern multiple cellular processes. A comprehensive understanding of the contexts in which specific dsRNAs bind specific proteins remains incomplete due, in part, to technical limitations. For example, the most commonly employed techniques to identify the repertoires of dsRNAs that bind a given dsRBP are limited by availability of specific antibodies for protein immunoprecipitation. Reciprocally, identification of the repertoires of proteins that bind a given dsRNA using IP-mass spec approaches is limited by an inability to reveal specific ‘reader domains’ that recognize the dsRNA of interest. In this study, we overcome these major technical challenges through the development and validation of dsRBP domain arrays and dsRBP-seq. Our discovery pipeline employs both techniques sequentially to identify new dsRNA-dsRBP interactions with domain-level resolution at high-throughput scale. We expect that this discovery pipeline can be applied widely across biological systems to enable high-throughput nomination of dsRNA-dsRBP candidate interactions for downstream biochemical validation.

Previous studies by our study team demonstrate the use of domain arrays as high-throughput approaches to discover specific reader domains that mediate protein-protein interactions. More recently, we demonstrate an extended utility of domain arrays to uncover interactions between domains containing RNA recognition motifs (RRMs) and poly ADP-ribose (PAR) chains of varying lengths. The chemical similarity of PAR chains to RNA established the rationale to further extend the use of RRM arrays, and establish the dsRBP array described in this study. Importantly, domains on the dsRBP array demonstrate selective preference for binding to dsRNA versus ssRNA substrates. Following validation, we sought to test the ability of the array to identify shared ‘readers’ of a given dsRNA of interest. To do so, we develop dsRBP-seq to identify putative dsRNA candidates to test on the dsRBP array. More specifically, we conduct dsRBP-seq to identify dsRNAs bound to catalytically inactive GST-tagged RNase III (GST-RNase III^E117A^). For technical feasibility and proof-of-principle, we select short dsRNA candidates for subsequent biochemical validation of dsRNA structure and RNase III binding. Following validation, we biotinylate *in vitro* transcribed dsRNA candidates, and subject the dsRNAs to the dsRBP array to discover putative domain binders. Using this approach, we discover and validate that RNase III-interacting dsRNAs that bind selectively to specific domains of other dsRNA binding proteins. Altogether, the technical advancement described here establishes a pipeline for dsRNA-dsRBP discovery. Future investigations of dsRBP-dsRNA interactions using this workflow can be started from either a candidate transcript or a candidate protein/domain. Domains can be cloned into a tag-expression vector and then for pulldowns with total RNA to isolate candidate dsRNAs of interest. Likewise, dsRNAs of interest can be biotinylated, and then applied to the dsRBP array to identify peptide ‘reader’ domains.

Our dsRBP array – dsRBP-seq pipeline identifies RNase III binding dsRNAs (*Sem1* and *Pfn1*) that also bind specific domains of Staufen 2. Staufen proteins contain multiple putative dsRNA binding domains. Here we highlight a domain specific dsRNA binding with Staufen 2 (STAU2). STAU2 is an RNA-binding protein with functions in mRNA localization, regulation of translation, and regulation of stauffen-mediated RNA decay (29). Previous studies suggest that all four dsRNA binding domains (dsRBDs) contribute towards STAU2 substrate RNA binding (30). Our in vitro pulldowns and dsRBP array suggest that SEM1 and PFN1 dsRNAs specifically bind STAU2 DRSM 3. Altogether, we demonstrate the ability of our discovery pipeline to nominate and validate domain-specific dsRNA-dsRBP interactions.

This current version of the dsRBP array establishes critical proof-of-principle and a foundation to incorporate innovations to address current technical limitations for this first version. For instance, we did not see strong binding of certain full-length proteins, such as MDA5 or RIG-I to poly (I:C). These limitations may be related to the immobilization of full-length proteins that may affect steric availability. Therefore, future arrays may benefit from testing modular domains instead of full-length proteins. Likewise, MDA5 exhibits strong substrate binding through the formation of filaments (31) which is not possible with immobilized protein monomers on the array. Conversely, technical constraints limit the range of dsRNA probes that can be tested on the array. Specifically, array probes are limited by length thresholds for in vitro transcription, and length limitations associated with end-labelling with biotin post transcription. Furthermore, some dsRNA-dsRBP interactions are sensitive to post-transcriptional modifications on the RNA substrate, as exemplified by RIG-I (32). In the absence of such modifications, recognition and binding of the unmodified dsRNA probes to a given dsRBD on the array might be limited. Furthermore, full-length proteins such as MDA5 exhibit different binding affinities to dsRNA *in vivo* versus *in vitro*. Therefore, future validation pipelines should include *in vivo* validation of putative interactions. Nevertheless, altogether the discovery pipeline reported in this study expands the technical capabilities to enable high throughput discovery of dsRNA-dsRBP interactions with domain level specificity across biological systems.

## ACKNOWLEDGEMENTS

C.A.I. acknowledges support from OCRA ECIG-2024-3-1467, NIH NCI R25CA181004, and UT MD Anderson Start Up Funds.

We would like to thank Dr. Yutong Sun and Dr. Chien-Chia Cheng for their help in generating the GST full length dsRNA sensors. We would like to thank Dr. Nakyung Oh, Ariana Paredes, and Tejas Patel for their help in assay development. We would like to thank Dr. Shannon Lauberth for guidance and protocols for in vitro transcription and biotin conjugation of the dsRNA probes. Finally, we would like to thank Dr. Sun Hur for providing protocols.

## AUTHOR CONTRIBUTIONS

N.Z. is responsible for experimental design, data generation, data analysis, original draft writing and manuscript preparation. L.S. is responsible for the experimental design, data generation, and analysis of the array experiments. B.C is responsible for data generation of the RIP seq experiment, and manuscript preparation. J.L. is responsible for developing the methodology, software generation, data curation for the dsRBP and RIP-seq analysis and IVT probe design. J.L. also assisted with original draft writing and manuscript preparation. I.J.M. generated the recombinant protein domains and the full length proteins used in the array, and assisted with recombinant protein purification in the GST-pulldown experiments. M.T.B. is responsible for the experimental design and data interpretation of the array experiments and IVT probe design. C.A.I is responsible for supervision of research, data interpretation, and manuscript preparation.

## DATA AVAILABILITY

Raw paired-end FASTQ files and strand-specific CPM-normalized bigWig coverage tracks have been deposited in the NCBI Gene Expression Omnibus; GSE348908 for the 9D5 RIP-seq series and GSE348907 for the GST–RNase III E117A dsRBP-seq series.

## SUPPLEMENTARY FIGURE LEGENDS

**Supplementary Figure S1: Layout of revised dsRNA binding domain array. (A)** Proteins and domains used in the array are listed for each block, where up to 12 domains or proteins were used. The center spot “C” of each block is GST alone as a negative control. All recombinant GST domains and proteins are arrayed in duplicate and arranged at different angles. Specific domains of the respective proteins are listed in brackets.

**Supplementary Figure S2: Determination and validation of dsRBP candidate Pfn1. (A)** GST-RNase III^E117A^ dsRBP-seq signal enrichment at the *Pfn1* locus. **(B)** 9D5 RIP-seq signal enrichment at the *Pfn1* locus. **(C)** Representative denaturing TBE-UREA gel of the in vitro transcribed (IVT) PFN1 candidate sequence. **(D)** Representative dot blot of poly(A), poly(I:C), and IVT PFN1 candidate RNA probed with anti-dsRNA (J2). Methylene blue staining shows total nucleic acids. **(E)** Representative dot blot showing dsRNA isolated after the GST-dsRBP pulldown.

**Supplementary Figure S3: High throughput discovery of Pfn1 binders. (A)** Representative SDS-PAGE gel of GST proteins used in pulldown. **(B)** Domain map of full-length STAU2. Double stranded RNA-binding motifs (DSRMs), microtubule binding domain (MTB) and C-terminal Staufen-swapping motif (ssm) are highlighted. **(C)** Representative fluorescence image from array version 2 of Cy3-detected IVT Pfn1. **(D)** Representative inset of block C, highlighting the signal from Pfn1. Full-length proteins incorporated on the block are listed with positive ‘hits’ highlighted based on signal from both technical replicates. **(E)** Representative blot showing dsRNA isolated after the GST-dsRBP pulldown using (IVT) Pfn1.

**Supplementary Table S1:** Information for all domains and proteins on the RBP array in Figure S1

**Supplementary Table S2:** Primer design for IVT transcripts in Figure 3E and Figure S2C

## REFERENCES

1. Hentze, M.W., Castello, A., Schwarzl, T. and Preiss, T. (2018) A brave new world of RNA-binding proteins. Nat Rev Mol Cell Biol, 19, 327–341.

2. Shi, L., Dasovich, M., Horton, J.R., Liu, H., Zhan, Q., Manickavinayaham, S., Jeter, C.R., Mixon, I.J., Zehrbach, N.M., Ishak, C.A. et al. (2025) Nuclear poly(A)-binding protein and nucleolin utilize their RNA recognition motifs to read PAR chains. Nucleic acids research, 53.

3. Ule, J., Jensen, K.B., Ruggiu, M., Mele, A., Ule, A. and Darnell, R.B. (2003) CLIP identifies Nova-regulated RNA networks in the brain. Science, 302, 1212–1215.

4. Ahmad, S., Mu, X., Yang, F., Greenwald, E., Park, J.W., Jacob, E., Zhang, C.Z. and Hur, S. (2018) Breaching Self-Tolerance to Alu Duplex RNA Underlies MDA5-Mediated Inflammation. Cell, 172, 797–810 e713.

5. Butter, F., Scheibe, M., Morl, M. and Mann, M. (2009) Unbiased RNA-protein interaction screen by quantitative proteomics. Proc Natl Acad Sci U S A, 106, 10626–10631.

6. Lin, H.Y., Xiang, J., Han, Y., Peng, X.X., Li, H. and Liu, X.J. (2026) Identification of poly(I:C) interacted proteins and their regulation in transcription level against bacterial infection in zebrafish. Front Immunol, 17, 1848091.

7. Hauk, P., Weeks, R. and Ostermeier, M. (2023) A CRISPR-dCas9 System for Assaying and Selecting for RNase III Activity In Vivo in Escherichia coli. CRISPR J, 6, 43–51.

8. Walton, J., Blagih, J., Ennis, D., Leung, E., Dowson, S., Farquharson, M., Tookman, L.A., Orange, C., Athineos, D., Mason, S. et al. (2016) CRISPR/Cas9-Mediated Trp53 and Brca2 Knockout to Generate Improved Murine Models of Ovarian High-Grade Serous Carcinoma. Cancer Res, 76, 6118–6129.

9. Bhola, M., Abe, K., Orozco, P., Rahnamoun, H., Avila-Lopez, P., Taylor, E., Muhammad, N., Liu, B., Patel, P., Marko, J.F. et al. (2024) RNA interacts with topoisomerase I to adjust DNA topology. Mol Cell, 84, 3192–3208 e3111.

10. Andrews, S. (2010), In Babraham Bioinformatics, Babraham Institute.

11. Ewels, P., Magnusson, M., Lundin, S. and Käller, M. (2016) MultiQC: summarize analysis results for multiple tools and samples in a single report. Bioinformatics, 32, 3047–3048.

12. Krueger, F., James, F., Ewels, P., Afyounian, E., Weinstein, M., Schuster-Boeckler, B. and Hulselmans, G. (2023). Zenodo.

13. Dobin, A., Davis, C.A., Schlesinger, F., Drenkow, J., Zaleski, C., Jha, S., Batut, P., Chaisson, M. and Gingeras, T.R. (2013) STAR: ultrafast universal RNA-seq aligner. Bioinformatics, 29, 15–21.

14. Danecek, P., Bonfield, J.K., Liddle, J., Marshall, J., Ohan, V., Pollard, M.O., Whitwham, A., Keane, T., McCarthy, S.A., Davies, R.M. et al. (2021) Twelve years of SAMtools and BCFtools. GigaScience, 10, giab008.

15. Ramírez, F., Ryan, D.P., Grüning, B., Bhardwaj, V., Kilpert, F., Richter, A.S., Heyne, S., Dündar, F. and Manke, T. (2016) deepTools2: a next generation web server for deep-sequencing data analysis. Nucleic Acids Research, 44, W160–W165.

16. Robinson, J.T., Thorvaldsdóttir, H., Winckler, W., Guttman, M., Lander, E.S., Getz, G. and Mesirov, J.P. (2011) Integrative genomics viewer. Nature Biotechnology, 29, 24–26.

17. Chen, Y., Chen, L., Lun, A.T.L., Baldoni, P.L. and Smyth, G.K. (2025) edgeR v4: powerful differential analysis of sequencing data with expanded functionality and improved support for small counts and larger datasets. Nucleic Acids Research, 53, gkaf018.

18. Espejo, A., Cote, J., Bednarek, A., Richard, S. and Bedford, M.T. (2002) A protein-domain microarray identifies novel protein-protein interactions. Biochem J, 367, 697–702.

19. Dasgupta, S., Fernandez, L., Kameyama, L., Inada, T., Nakamura, Y., Pappas, A. and Court, D.L. (1998) Genetic uncoupling of the dsRNA-binding and RNA cleavage activities of the Escherichia coli endoribonuclease RNase III--the effect of dsRNA binding on gene expression. Mol Microbiol, 28, 629–640.

20. Blaszczyk, J., Gan, J., Tropea, J.E., Court, D.L., Waugh, D.S. and Ji, X. (2004) Noncatalytic assembly of ribonuclease III with double-stranded RNA. Structure, 12, 457–466.

21. Sun, W., Pertzev, A. and Nicholson, A.W. (2005) Catalytic mechanism of Escherichia coli ribonuclease III: kinetic and inhibitor evidence for the involvement of two magnesium ions in RNA phosphodiester hydrolysis. Nucleic Acids Res, 33, 807–815.

22. Inada, T., Kawakami, K., Chen, S.M., Takiff, H.E., Court, D.L. and Nakamura, Y. (1989) Temperature-sensitive lethal mutant of era, a G protein in Escherichia coli. J Bacteriol, 171, 5017–5024.

23. Blaszczyk, J., Tropea, J.E., Bubunenko, M., Routzahn, K.M., Waugh, D.S., Court, D.L. and Ji, X. (2001) Crystallographic and modeling studies of RNase III suggest a mechanism for double-stranded RNA cleavage. Structure, 9, 1225–1236.

24. Ishak, C.A., Marhon, S.A., Tchrakian, N., Hodgson, A., Loo Yau, H., Gonzaga, I.M., Peralta, M., Lungu, I.M., Gomez, S., Liang, S.B. et al. (2025) Chronic Viral Mimicry Induction following p53 Loss Promotes Immune Evasion. Cancer Discov, 15, 793–817.

25. Bou-Nader, C., Juma, K.M., Bothra, A., Brasington, A.J., Ghirlando, R., Suzuki, M., Garboczi, D.N., Leppla, S.H. and Zhang, J. (2025) Structural basis of double-stranded RNA recognition by the J2 monoclonal antibody. Nat Commun, 17, 635.

26. Murayama, T., Nakayama, J., Jiang, X., Miyata, K., Morris, A.D., Cai, K.Q., Prasad, R.M., Ma, X., Efimov, A., Belani, N. et al. (2024) Targeting DHX9 Triggers Tumor-Intrinsic Interferon Response and Replication Stress in Small Cell Lung Cancer. Cancer Discov, 14, 468–491.

27. Schonborn, J., Oberstrass, J., Breyel, E., Tittgen, J., Schumacher, J. and Lukacs, N. (1991) Monoclonal antibodies to double-stranded RNA as probes of RNA structure in crude nucleic acid extracts. Nucleic Acids Res, 19, 2993–3000.

28. Kitagawa, Y., Matsumoto, T., Okuhara, E. and Shikata, E. (1977) Immunogenicity of rice dwarf virus-ribonucleic acid. Tohoku J Exp Med, 122, 337–343.

29. Park, E., Gleghorn, M.L. and Maquat, L.E. (2013) Staufen2 functions in Staufen1-mediated mRNA decay by binding to itself and its paralog and promoting UPF1 helicase but not ATPase activity. Proc Natl Acad Sci U S A, 110, 405–412.

30. Heber, S., Gáspár, I., Tants, J.N., Günther, J., Moya, S.M.F., Janowski, R., Ephrussi, A., Sattler, M. and Niessing, D. (2019) Staufen2-mediated RNA recognition and localization requires combinatorial action of multiple domains. Nat Commun, 10, 1659.

31. Ahmad, S., Mu, X., Yang, F., Greenwald, E., Park, J.W., Jacob, E., Zhang, C.Z. and Hur, S. (2018) Breaching Self-Tolerance to Alu Duplex RNA Underlies MDA5-Mediated Inflammation. Cell, 172, 797–810.e713.

32. Hornung, V., Ellegast, J., Kim, S., Brzózka, K., Jung, A., Kato, H., Poeck, H., Akira, S., Conzelmann, K.K., Schlee, M. et al. (2006) 5’-Triphosphate RNA is the ligand for RIG-I. Science, 314, 994–997.

