## Supplementary material for "Development of a high-throughput protein array to discover selective dsRNA binding domain interactions": Figure_S1

### SUPPLEMENTARY FIGURE S1

**A**

#### dsRNA Binding Domain Array Version 2

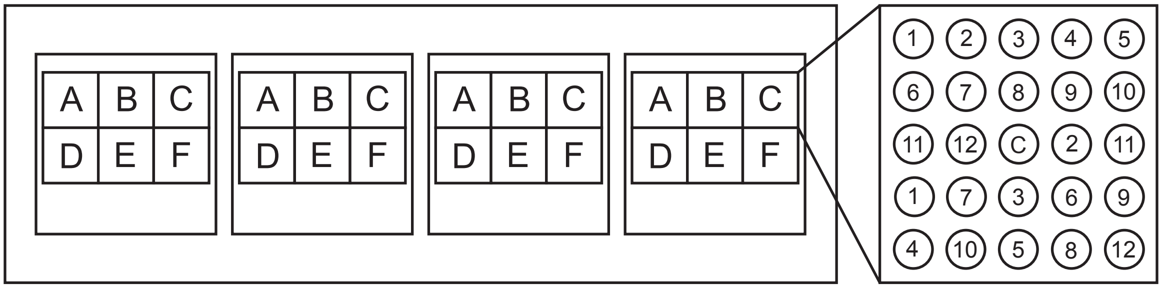


**Block A**

DSRM
1. ADAD1 DSRM
2. ADAD2 DSRM
3. ADAR DSRM(1)
4. ADAR DSRM(2)
5. ADAR DSRM(3)
6. ADARB2 DSRM(1)
7. ADARB2 DSRM(2)
8. DDX9/DHX9 DSRM(1)
9. DDX9/DHX9 DSRM(2)
10. DGCR8 DSRM(1-2)
11. DROSHA DSRM
12. DUS2L DSRM

**Block B**

DSRM
1. Kanadaptin DRBM
2. MRPL44 DSRM
3. NFAT90/ILF3 DSRM(1)
4. NFAT90/ILF3 DSRM(2)
5. PACT DSRM(1-3)
6. PKR DSRM(1-2)
7. RED1/ADARB1 DSRM(1)
8. RED1/ADARB1 DSRM(2)
9. STAU1 DSRM(1)
10. STAU1 DSRM(2)
11. STAU1 DSRM(3)
12. STAU2 DSRM(1-2)

**Block C**

DSRM
1. STAU2 DSRM(3)
2. STAU2 DSRM(4)
3. STRBP DSRM(1)
4. STRBP DSRM(2)
5. TRBP DSRM(1)
6. TRBP DSRM(2)
7. TRBP DSRM(3)
8. DHX30 DRBM
9. DHX36 DRBM

**Block D**

DND1/DSRM
1. ACF DND1_DSRM
2. DND1 DND1_DSRM
3. MGC27016 DND1_DSRM
4. SON DND1_DSRM

Helicase

5. MDA5 Helicase
6. RIG_I Helicase
7. LGP2 Helicase

RIG-I-C

8. MDA5 RIG-I_C
9. RIG_I RIG-I_C
10. LGP2 RIG-I_C

LRR

11. TLR3 LRR12_20

**Block E**

DSRI
1. OASL DSRI
2. OAS1 DSRI
3. OAS2 DSRI
4. OAS3 DSRI

OB Fold

5. DHX30 OB Fold
6. DHX15 OB Fold
7. DHX36 OB Fold

**Block F**

Full Length
1. MDA5 FL
2. MDA5 $\Delta$2CARD
3. RIG-I
4. LGP2
5. TLR3
6. RNase III WT
7. RNase III E117Q
8. RNase III E117A
9. RNase III E117K
10. RNase III D45A
11. RNase H
