## Supplementary figures and images for "Development of a high-throughput protein array to discover selective dsRNA binding domain interactions"

### Figure_S2

#### SUPPLEMENTARY FIGURE S2

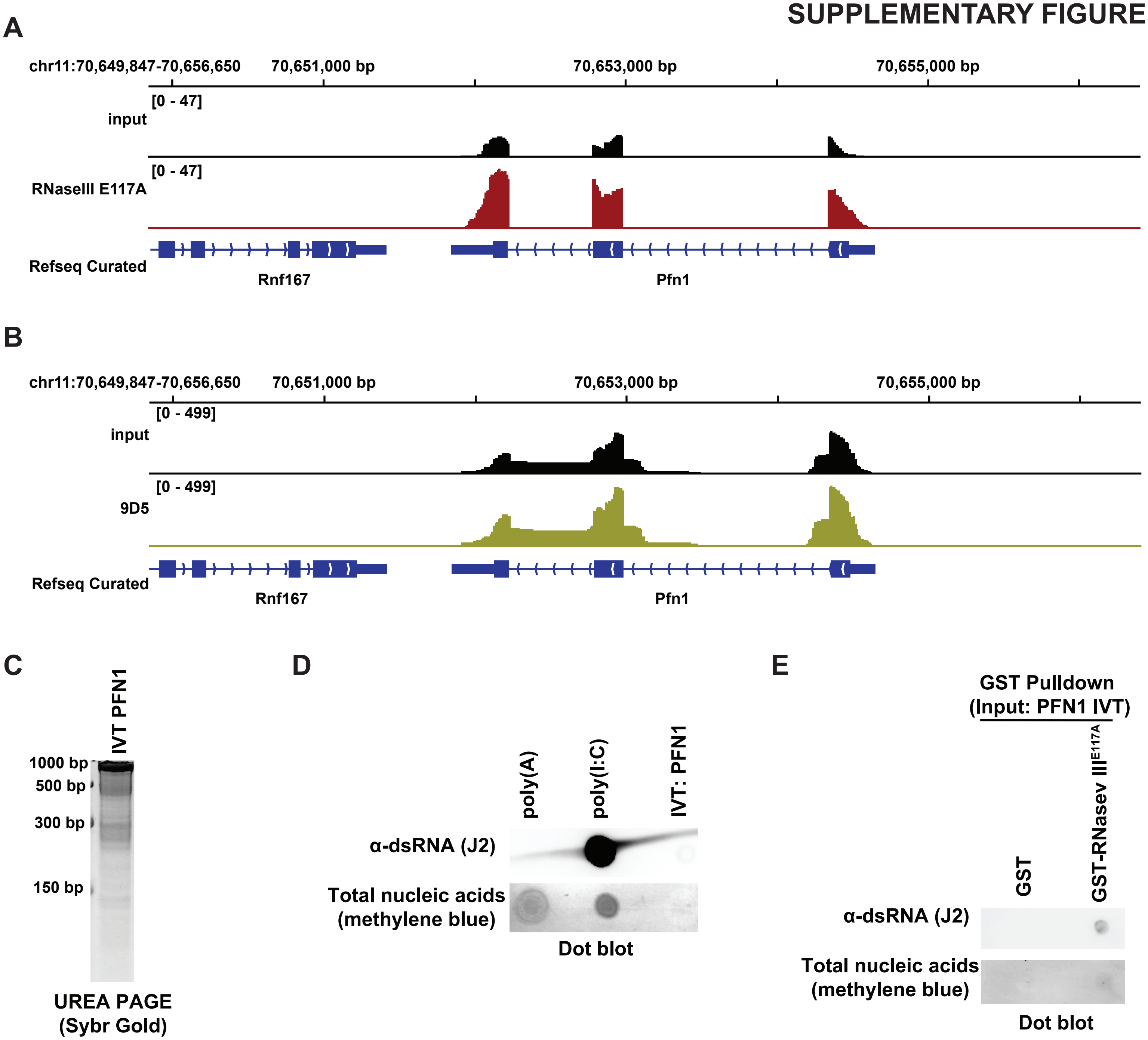
