## Supplementary material for "Development of a high-throughput protein array to discover selective dsRNA binding domain interactions": Figure_S3

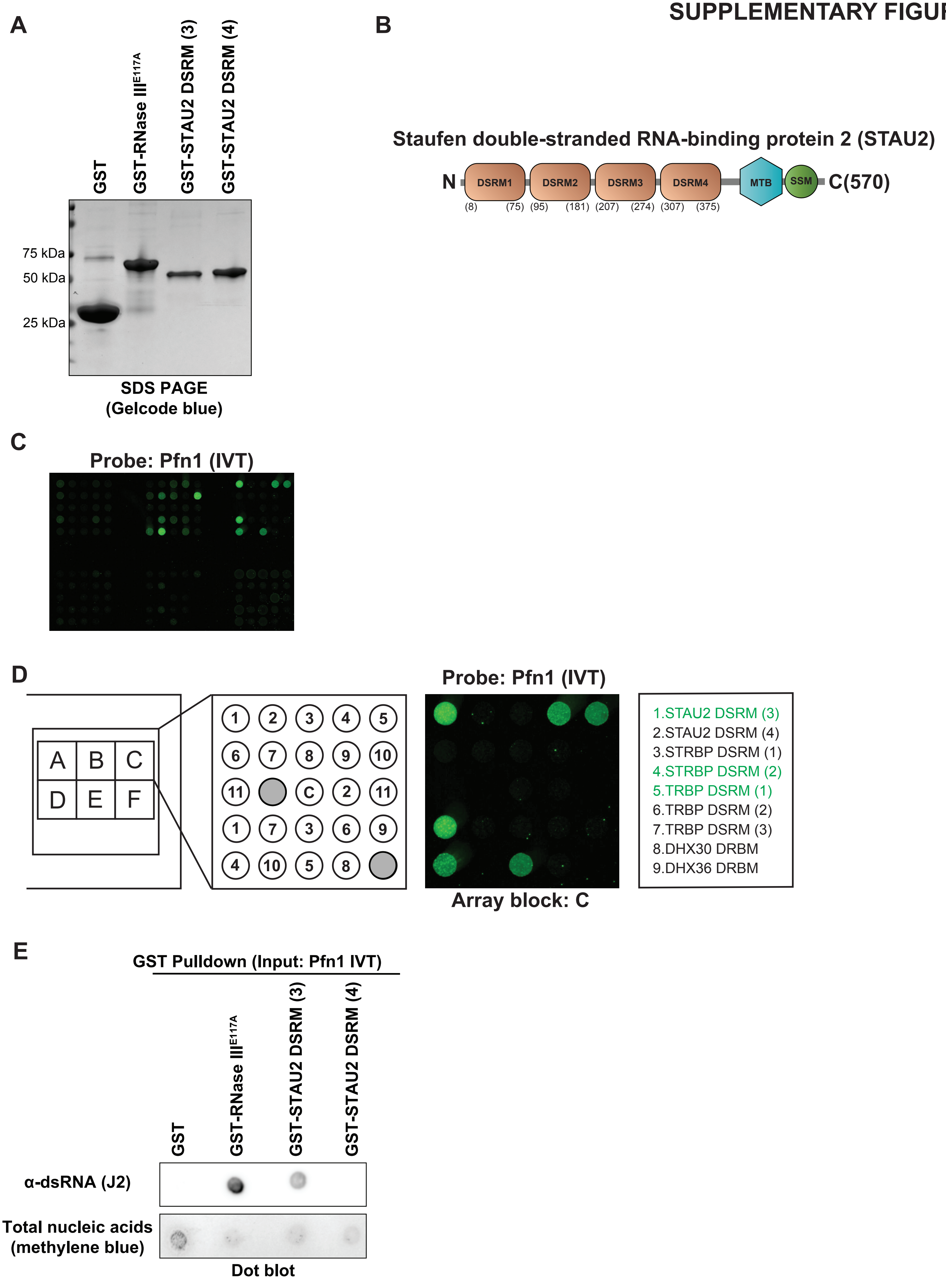
A
GST
GST-RNase III$^{E117A}$
GST-STAU2 DSRM (3)
GST-STAU2 DSRM (4)
75 kDa
50 kDa
25 kDa
SDS PAGE
(Gelcode blue)
B
Staufen double-stranded RNA-binding protein 2 (STAU2)
N
DSRM1
DSRM2
DSRM3
DSRM4
MTB
SSM
C(570)
(8) (75) (95) (181) (207) (274) (307) (375)
C
Probe: Pfn1 (IVT)
D
A B C
D E F
1 2 3 4 5
6 7 8 9 10
11 C 2 11
1 7 3 6 9
4 10 5 8
Probe: Pfn1 (IVT)
Array block: C
1.STAU2 DSRM (3)
2.STAU2 DSRM (4)
3.STRBP DSRM (1)
4.STRBP DSRM (2)
5.TRBP DSRM (1)
6.TRBP DSRM (2)
7.TRBP DSRM (3)
8.DHX30 DRBM
9.DHX36 DRBM
E
GST Pulldown (Input: Pfn1 IVT)
GST
GST-RNase III$^{E117A}$
GST-STAU2 DSRM (3)
GST-STAU2 DSRM (4)
α-dsRNA (J2)
Total nucleic acids
(methylene blue)
Dot blot
